# Evidence for the Emergence of β-Trefoils by ‘Peptide Budding’ from an IgG-like β-Sandwich

**DOI:** 10.1101/2021.10.04.462989

**Authors:** Liam M. Longo, Rachel Kolodny, Shawn E. McGlynn

## Abstract

As sequence and structure comparison algorithms gain sensitivity, the intrinsic interconnectedness of the protein universe has become increasingly apparent. Despite this general trend, β-trefoils have emerged as an uncommon counterexample: They are an isolated protein lineage for which few, if any, sequence or structure associations to other lineages have been identified. If β-trefoils are, in fact, remote islands in sequence-structure space, it implies that the oligomerizing peptide that founded the β-trefoil lineage itself arose *de novo*. To better understand β-trefoil evolution, and to probe the limits of fragment sharing across the protein universe, we identified both ‘β-trefoil bridging themes’ (evolutionarily-related sequence segments) and ‘β-trefoil-like motifs’ (structure motifs with a hallmark feature of the β-trefoil architecture) in multiple, ostensibly unrelated, protein lineages. The success of the present approach stems, in part, from considering β-trefoil sequence segments or structure motifs rather than the β-trefoil architecture as a whole, as has been done previously. The newly uncovered inter-lineage connections presented here suggest a novel hypothesis about the origins of the β-trefoil fold itself – namely, that it is a derived fold formed by ‘budding’ from an Immunoglobulin-like β-sandwich protein. These results demonstrate how the emergence of a folded domain from a peptide need not be a signature of antiquity and underpin an emerging truth: few protein lineages escape nature’s sewing table.

## Introduction

Proteins are often approximated as discrete domains from distinct evolutionary lineages. Such classification is powerful, foremost because it can serve as a framework for understanding how proteins within a family change over time, but also because it naturally lends itself to naming and thus aids in communication (as is true for taxonomy in general). Nevertheless, this notion of separability is, ultimately, an approximation: Regardless of whether structure ^1^, sequence^2–5^, or both structure and sequence^6,7^ are considered, evolutionarily related segments between proteins that lack global sequence similarity are detectable and common. These studies emphasize the interconnected, ‘patchwork’ nature of protein evolution and reveal that proteins might be best understood as being comprised of multiple segments, each with its own independent evolutionary history or structural properties. In extreme cases, a protein domain is not unlike the genome that encodes it: a composite of elements that evolves by accretion, exchange, and loss. When these segments are highly conserved within diverse protein families, or overlap with important functional sites, they can provide insight into early protein evolution^7^ and reveal distant evolutionary relationships^8^.

One protein lineage of particular interest is the β-trefoil (ECOD X-group 6), which is characterized by a common ancestor^9^ and a pseudo-three-fold axis of rotational symmetry. Although originally proposed to be related to EGF (X-group 389) and ecotin (X-group 521) by gene duplication and fusion^10^, proteome-wide sequence analyses^2,3,6^ and an experimental fragmentation study^11^ have found no support for this hypothesis; instead, the ancestral state of the β-trefoil was most likely a trimerizing peptide that recapitulated the β-trefoil fold. Given that β-trefoils seem to comprise a rare island in an otherwise highly-connected sequence-structure landscape, the origins of the precursor β-trefoil peptide are unclear and potentially the result of a *de novo* emergence event.

Recently, Tenorio and coworkers proposed a link between the β-trefoil and the β-propellor (X-group 5) on the basis of *ab initio* folding simulations^12^, in which a four-stranded β-trefoil motif was predicted to adopt a β-propellor blade-like structure by the Robetta protein structure prediction server^13^. Although the putative evolutionary relationship between β-propellors and β-trefoils was not validated by a detailed sequence analysis, this study nevertheless provided two valuable insights: First, the elements under consideration were β-trefoil subdomains, not the entire β-trefoil architecture; and second, the subdomains themselves were predicted to be metamorphic (*i*.*e*., adopt different structures in different contexts). Finally, Tenorio and coworkers reignited the >20-year-old debate^10,12^ about whether β-trefoils are related to, and perhaps derived from, another common β-protein architecture.

We now report a systematic search for β-trefoil-like sequence segments (bridging themes^8^) and structure motifs (β-trefoil-like motifs) across the known protein universe, foremost to understand the evolutionary history of β-trefoils but also as a test for the ‘patchwork model’ of protein evolution. Whereas previous searches have generally considered the β-trefoil architecture in its entirety, we now take a segment-centric approach^14^ for improved sensitivity. We observe convergently evolved β-trefoil-like structural motifs in two unrelated protein lineages and identify 68 cases of statistically significant (E-value ≤ 1×10^−3^) sequence segment sharing between a β-trefoil protein and a *globally* unrelated protein lineage. Taken together, these results demonstrate that the β-trefoil is nowhere near as isolated as previously thought; instead, β-trefoil proteins appear to be a reservoir for sequence innovation in other protein lineages and *vice versa*. The implications of these results with respect to the emergence of the founding β-trefoil are discussed, and we argue that β-trefoils – a relatively recent protein architecture innovation^9,15–17^ – are potentially derived from a sequence segment that was once embedded in a different β-protein architecture, specifically, the much more ancient^15–17^ Immunoglobulin-like β-sandwich (IgG-like β-sandwich; X-group 11). This result is a reminder that emergence from a peptide need not necessarily be taken as a signature of great age. Ultimately, as our analytical tools become more and more sensitive, it is becoming increasingly clear that, to rephrase John Donne, no protein lineage is an island entire of itself.

## Methods

### Bridging Theme Search

Bridging themes are sequence segments that link, or bridge, two protein domains from different evolutionary lineages that are presumed to be unrelated based on global sequence and structure considerations^8,18^. For the purposes of this report, a protein evolutionary lineage is defined as the X-group level of hierarchy in the Evolutionary Classification of Domains (ECOD) database^19–21^. Within an ECOD X-group, domains most often share the same global architecture and are hypothesized to have emerged from a common origin (even if more evidence is needed to establish an unambiguous evolutionary relationship). Different X-groups are classified as entirely distinct evolutionary lineages and so a bridging theme is defined here as an instance of sequence segment sharing between two different ECOD X-groups.

To build a dataset of HMMs modeling structurally relevant parts of the β-trefoil domains, we employed the following procedure: First, we built multiple sequence alignments (MSAs) from the 128 β-trefoil domains in ECOD 70% NR (X-group 6; version develop271) by searching Uniclust3021 with HHblits^23^. For each domain we calculated the residue ranges corresponding to each β-strand with the STRIDE algorithm^24^. Using a sliding window of 4 β-strands (*e*.*g*., β-strands 1-4, 2-5 and so on), which is the length of the repetitive structural element that comprises the β-trefoil architecture, we restricted the MSAs and built hidden Markov models (HMM) using hmmbuild from the HMMER package^25,26^. Finally, we searched for similarities to all ECOD domains using hmmsearch from the HMMER package. Among the similarities found, we selected the ones with E-values < 1×10^−3^ and not from X-group 6 (the β-trefoils) for further analysis.

To further confirm the statistical significance of the bridging themes themselves, we shuffled and realigned the two associated sequence segments, fit the resulting alignment score to an extreme value distribution, and estimated the likelihood of the observed alignment. In the subsequent text and figures, p-value refers to this analysis whereas E-values are taken from the HMMER output. The bridging themes identified in this work are reported in **Supplemental File 1**.

### β-Trefoil-like (βTL) Motif Search

The structural motif used to query the Protein Databank (PDB) was extracted from the sequence-symmetrized β-trefoil Phifoil, which is derived from the folding nucleus of human fibroblast growth factor-1 (FGF-1) and adopts an idealized structure^27^. Specifically, residues 52-60 and 77-87 from ECOD domain e4ow4A1 were used for the search motif. The rationale behind this choice of motif is discussed in greater detail in the section β**-Trefoil-like Motifs Occur in Two Ancient** β**-Protein Lineages**, and relates to the rarity of this motif and its essentiality to β-trefoil structure and folding^28^. The search motif was aligned against all domains in the ECOD F99 database (version develop275) with TM-align^29^. TM-scores were normalized to the length of search motif, which is constant across all alignments. The top hits (excluding β-trefoils) were inspected visually. Alignments to the search motif yielded TM-scores of 0.52 for e2×9wA3 and 0.53 for e2o1cA1. A TM-score > 0.5 is indicative of statistically significant structural similarity^30^. The alignment outputs are included as **Supplemental File 2**.

## Results

### β-Trefoils have Bridging Themes with up to 68 X-Groups, Foremost IgG-like β-Sandwiches

In total, 68 different evolutionary lineages share at least one bridging theme with a β-trefoil, comprising just under 3% of all evolutionary lineages in ECOD (**Figure 1A**). Unsurprisingly, the greatest number of shared themes is with other β-proteins, though the themes themselves are largely metamorphic (*i*.*e*., they adopt different conformations). Among the X-groups that share a theme with the β-trefoil are EGF (first suggested by Mukhopadhyay^10^) and the β-propeller (first suggested by Tenorio and coworkers^12^), with 8 and 7 unique themes, respectively (E-value < 1×10^−3^; **Figure 1B** and **Supplemental File 1**). The evolutionary lineage with by far the greatest number of unique bridging themes to the β-trefoil, however, was the IgG-like β-sandwich, with 48 themes in total. Increasing the stringency of the theme cutoff does not change the result that IgG-like β-sandwiches are the most connected evolutionary lineage to the β-trefoil (**Figure 1B**).

**Figure 1.**
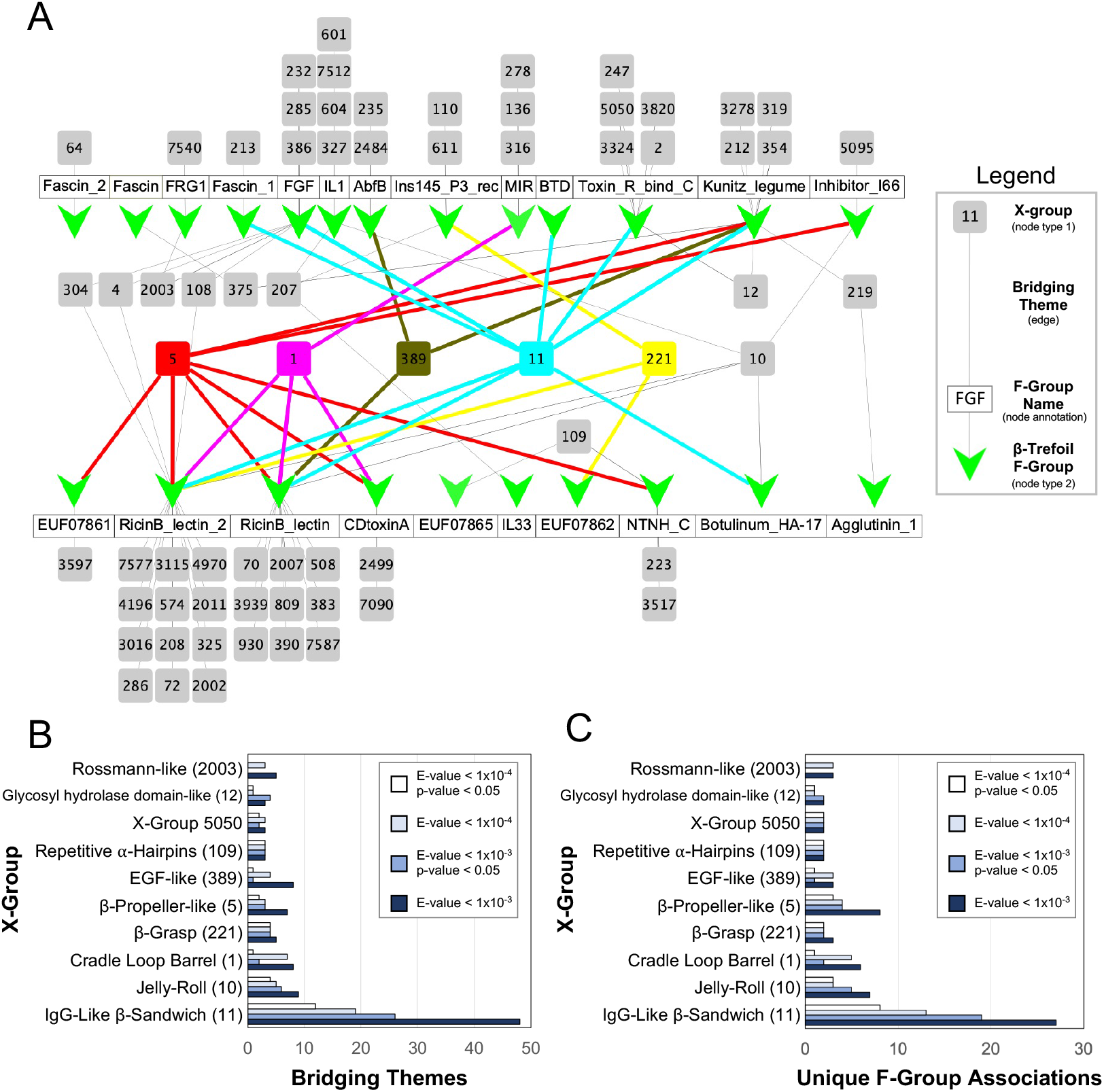
Bridging themes to the β-trefoil evolutionary lineage. **A**. Network of bridging themes (edges) between 23 β-trefoil families (nodes represented by green Vs, defined as an F-group) and 68 other evolutionary lineages (nodes represented by grey and colored boxes, defined as an X-group). The network was constructed using an E-value cutoff of 1×10^−3^. For clarity, evolutionary lineages with a bridging theme to only one β-trefoil family are tiled above or below the name of the associated family (white boxes). Nodes associated with evolutionary lineages of special note are colored as follows: X-group 5 (β-propeller) is red; X-group 1 (cradle loop barrel) in magenta; X-group 389 (EGF) in olive; X-group 11 (IgG-like β-sandwich) in cyan; and X-group 221 (β-grasp) in yellow. The IgG-like β-sandwich is associated with the most β-trefoil families (8 in total). The 7 β-trefoil families for which no bridging themes were found are not shown. The network graph was rendered with Cytoscape (cytoscape.org) **B**. Bridging theme counts between the β-trefoil and other evolutionary lineages. IgG-like β-sandwiches are associated with the greatest number of bridging themes to the β-trefoil lineage, even with increasingly stringent statistical cutoffs (see **Methods** for more details). Note that there can be, and often are, multiple, unique bridging themes within a β-trefoil F-group to the same X-group (*i*.*e*., an edge in the network can be associated with more than one shared sequence segment). For clarity, only the top 10 most connected evolutionary lineages are shown. **C**. The number of unique pairwise F-group associations between the β-trefoil and another protein lineage. For clarity, only the top 10 most connected evolutionary lineages are shown.

Of the 32 known β-trefoil families (*i*.*e*., ECOD F-groups), 23 have at least one bridging theme to a different evolutionary lineage, with the IgG-like β-sandwich linked to the greatest number of β-trefoil families, 8 in total (E-value > 1×10^−3^; **Figure 1A**). Although β-propellers have a similar number of associated β-trefoil families with 7 (**Figure 1A**), each of these associations is the result of fewer unique sequence themes than with the IgG-like β-sandwich – an average of 1 *vs*. 6, respectively – indicating a weaker evolutionary association. Likewise, the number of unique F-group to F-group associations between the β-trefoil and the IgG-like β-sandwich is greater than with any other X-group.

### The Phylogenetic Distribution of Bridging Themes Suggests that an Early β-Trefoil is related to an IgG-like β-Sandwich

The distribution of bridging themes across the phylogenetic trees of both protein lineages may help clarify the nature of their evolutionary relationship. If bridging themes are widely distributed across *both phylogenetic trees*, it would be an indication of either i) significant sequence sharing early in the evolution of both folds, and perhaps a shared evolutionary origin; or ii) a history of repeated, independent fragment sharing events. In cases where one evolutionary lineage is much older than the other, the second interpretation (repeated sharing) is more consistent with the data. However, if bridging themes are widely distributed across the phylogenetic tree of the younger fold but narrowly distributed across the phylogenetic tree of the older fold, it suggests that a sequence sharing event happened very early in the evolutionary history of the younger fold, perhaps corresponding to an emergence event. Finally, if the distribution of bridging themes is narrow in both phylogenetic trees, it suggests a more recent theme sharing event, unrelated to the early evolution of either fold.

The phylogenetic distribution of theme sharing between the IgG-like β-sandwich and the β-trefoil more closely follows the second case above (**Figure 2A**, left panel): A phylogenetically wide distribution of β-trefoil F-groups are associated with just a single H-group (that is, a group of F-groups; specifically, 11.1.1) of IgG-like β-sandwiches, an evolutionary lineage with 47 H-groups in total. Contrast this phylogenetic distribution with that of the β-propeller bridging themes (**Figure 2A**, right panel), the evolutionary lineage with the second most associations to β-trefoil F-groups. In this case, the most prominent H-group (5.1.4) is associated with just four β-trefoil F-groups, and these F-groups are more closely related to each other than those associated with the IgG-like β-sandwich. For another example, see the distribution of β-grasp bridging themes (**Figure S1**). Indeed, the IgG-like β-sandwich is unique in the breadth of associations across the β-trefoil phylogenetic tree that is achieved by just a single H-group. However, it should be noted that within the IgG-like β-sandwich evolutionary lineage, X-group 11.1.1 is the largest and most sequence diverse. The fact that detectable signals are not present in all β-trefoil families may relate to the complex evolution of this lineage29 or to the fact that β-trefoils generally have highly diverged sequences^9^.

**Figure 2.**
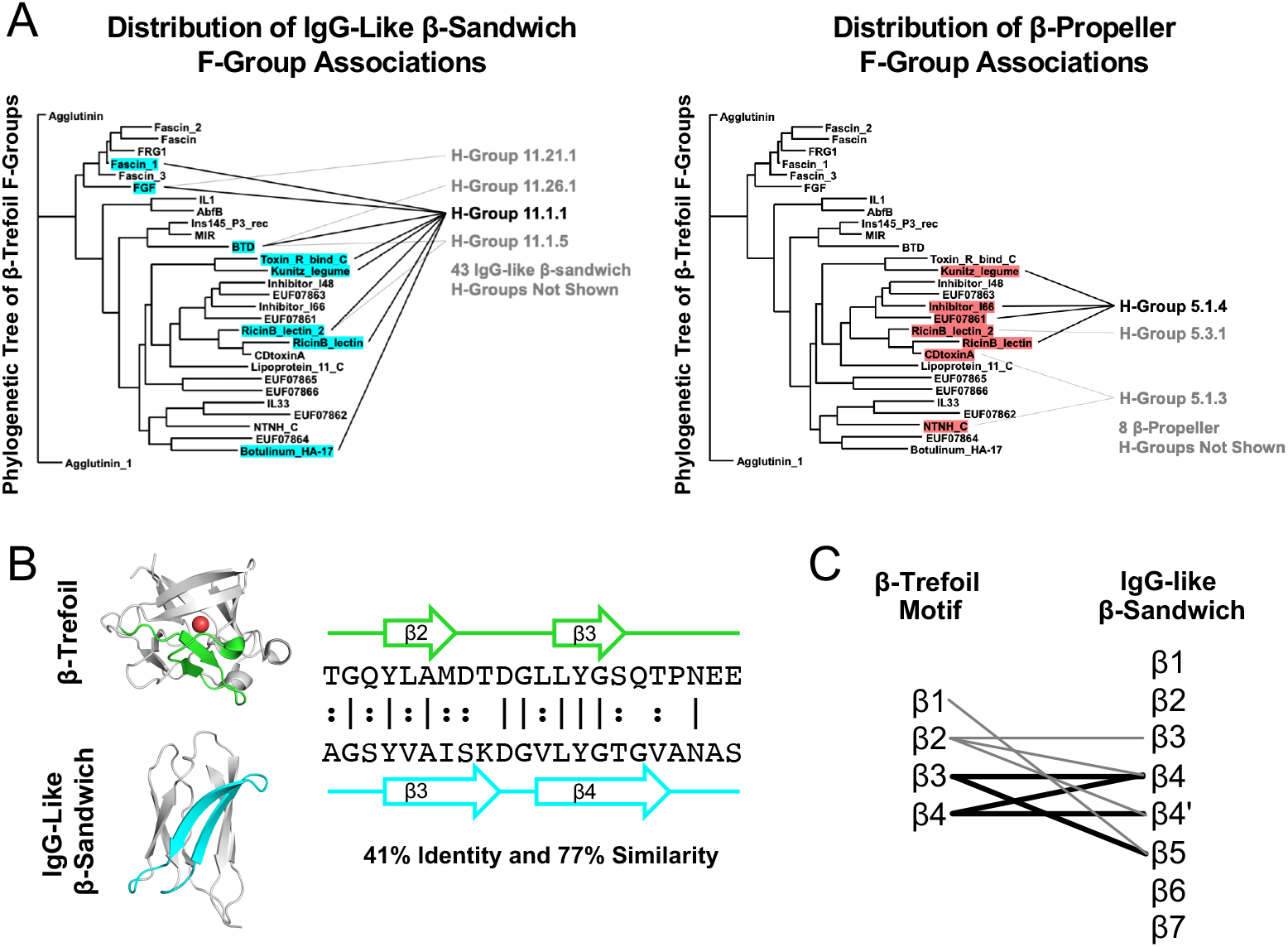
The IgG-like β-sandwich as a candidate progenitor of the β-trefoil. **A**. The distribution of bridging themes across the β-trefoil phylogenetic tree. β-trefoil families with a bridging theme are highlighted following the color scheme in **Figure 1**. For the IgG-like β-sandwich, the specific IgG-like β-sandwich H-group (*i*.*e*., the group of F-groups) that is associated with the β-trefoil family is noted. The IgG-like β-sandwich is not only associated with the most β-trefoil families, these families are well-distributed across the β-trefoil evolutionary tree (unlike, for example, the β-propeller). And, while the bridging themes of other protein lineages span multiple H-groups, all of the β-trefoil families associated with the IgG-like β-sandwich have at least one bridging theme to the same H-group, 11.1.1, which is just one out of 47 H-groups associated with this X-group (though it is the also largest and most diverse one). The β-trefoil phylogenetic tree was adapted from ECOD (http://prodata.swmed.edu/ecod/complete/famtree?tid=6.1.1). **B**. An example bridging theme between a β-trefoil and an IgG-like β-sandwich, ECOD domains e1rg8A1 and e4rbmA3, respectively. By aligning the themes and denoting the positions of the β-strands, strand associations can be generated. Structure figures were rendered in PyMOL (pymol.org). **C**. Strand associations between the β-trefoil and the IgG-like β-sandwich for themes with an E-value < 1×10^−4^ that have a structure model for both sequence segments. Although localized to β3-β5 in the IgG-like β-sandwich structure, the specific strand associations are unclear, perhaps due to rearrangement events in either the β-trefoil and/or IgG-like β-sandwich lineages. Nevertheless, all four β-strands in the β-trefoil repetitive motif have some association to IgG-like β-sandwich. Thin lines correspond to a single bridging theme with this β-strand mapping whereas thick lines correspond to two themes.

In cases where the sequence associated with the bridging theme was present in the structure models of both domains, strand associations between the IgG-like β-propellor and the β-trefoil were identified to determine if they are consistent across the bridging themes (**Figure 2B** and **2C**). Although the strand associations implied by the theme alignments are not internally consistent – that is, there is no simple one-to-one correspondence between strands in the IgG-like sandwich and strands in the β-trefoil – they are localized to a specific region of the β-sandwich structure and, between them, encompass all four β-strands of the β-trefoil (the structure of which is described in greater detail the next section). Although none of the identified themes adopt a β-trefoil like structure within an IgG-like β-sandwich domain, perhaps such a structure has arisen *de novo* – thereby demonstrating the structural potential of the IgG-like β-sandwich to host a β-trefoil-like motif.

### β-Trefoil-like Structure Motifs Occur in Just Two Ancient β-Protein Lineages, Including the IgG-like β-Sandwich

The β-trefoil architecture is comprised of three symmetrically juxtaposed structural subdomains referred to as β-trefoil motifs. Each β-trefoil motif is made up of four β-strands and an absolutely conserved buried water molecule^28^ (**Figure 3**). The conserved water molecule forms hydrogen bonds to three distinct backbone sites, two β-strands and a loop, to form a hallmark structural feature that fundamentally enables the β-trefoil motif and architecture^28^. To search for β-trefoil-like motifs in other protein lineages, we focused on the conserved water molecule and the structural elements that interact with it, namely β-strands β1, β2, and the loop after β3 (canonical β-trefoil motif strand numbering). We define this structural motif as the minimal definition of the β-trefoil-like motif (βTL) because, unlike a simple β-hairpin, we expect this feature to be rare, if not unique to the β-trefoil fold altogether. To reduce family-specific bias, the structure used for searching was extracted from an idealized β-trefoil protein with fully symmetrized sequence and structure^32^ (see the **Methods** for more details).

**Figure 3.**
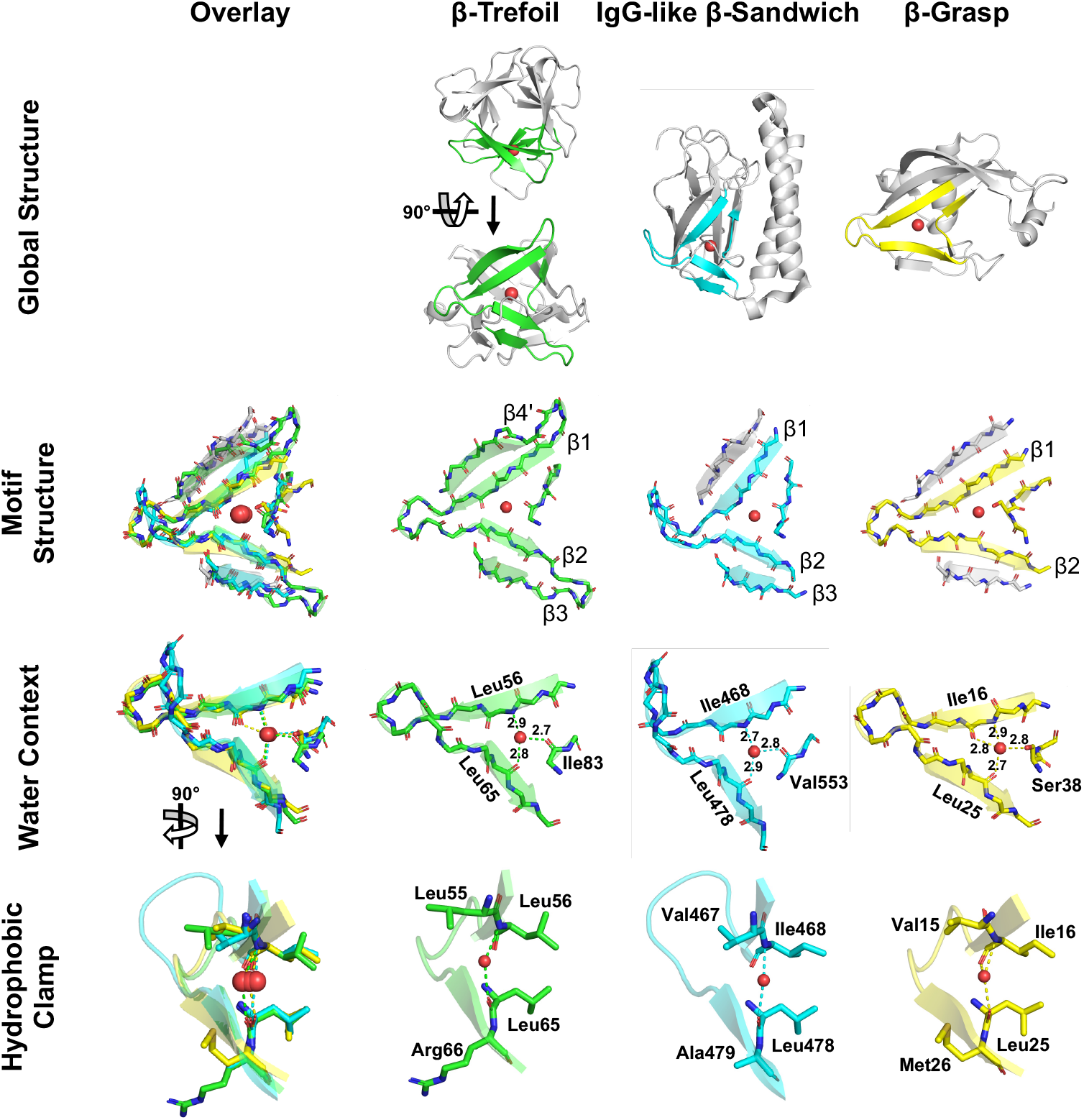
β-trefoil-like (βTL) motifs across the protein universe. The canonical β-trefoil motif (second column, green) comprises four β-strands, their intervening loops and turns, and a conserved water molecule. Shown here is a circular permutation of the canonical motif definition, comprised of β4’, β1, β2, and β3 and the subsequent loop, where β4’ refers to the final β-strand (β4) of the previous β-trefoil motif. Water molecules are indicated by a red sphere. β-strands shaded grey in the **Motif Structure** panel are surrogate β-strands, donated by other parts of the host domain, and are not part of the βTL motif proper but satisfy the same role as a canonical β-strand. Distances in the **Water Context** row are shown in Angstroms (Å). The structure motifs shown here were extracted from the following ECOD domains: β-trefoil: e3o4bA1 (F-group 6.1.1.8); IgG-like β-sandwich: e2×9wA3 (F-group 11.1.4.227); β-grasp: e2o1cA1 (F-group 221.4.1.3).

Structural alignments against the representative domains for each lineage of the ECOD database identified two independent protein lineages bearing a βTL motif (**Figure 3**). In both cases, the similarity of the identified βTL motif to a *bona fide* β-trefoil motif extends beyond the search motif itself to include β3 or β4’ (the last β-strand, β4, of the preceding β-trefoil motif, which packs against β1; see **Figure 3 Motif Structure**). Unlike the β-trefoil architecture, which is rotationally symmetric, the detected βTL motifs were found in two asymmetric architectures – the β-grasp (ECOD X-group 221) and the IgG-like β-sandwich. Despite divergent structural contexts, all of the βTL motifs adopt a highly similar conformation (**Figure 3 Motif Structure**) and conserved water position (**Figure 3 Water Context**). In each case, the residues on β1 and β2 that form a backbone hydrogen bond with the conserved water also form a ‘hydrophobic clamp’ with their sidechains (**Figure 3 Hydrophobic Clamp**). Although both evolutionary lineages associated with a β-trefoil-like motif also share bridging themes with β-trefoils, these bridging themes were not associated with either of the entire βTL motifs identified here, though for the β-grasp, there is evidence of partial overlap between a bridging theme and a β-trefoil-like motif. In addition, for the IgG-like β-sandwich, some sequence similarity to an extant β-trefoil is apparent. The novel βTL motifs are now described in turn.

*IgG-like* β*-sandwich*. The occurrence of a βTL motif in the IgG-like β-sandwich evolutionary lineage was identified in ECOD F-group 11.1.4.227 – a different H-group than that associated with the bridging themes (11.1.1). The fact that structure comparisons and sequence comparisons identified different protein elements is consistent with the observation that the shared themes we identified are mostly metamorphic (*i*.*e*., adopt different conformations in in different contexts). As βTL motifs were not detected in closely related F-groups, this motif appears to be a relatively recent, niche innovation. As in the canonical β-trefoil motif, the IgG-like β-sandwich βTL motif is characterized by a conserved water bound by three short (<3.0 Å) hydrogen bonds to the protein backbone. The IgG-like β-sandwich βTL motif comprises β1, β2, β3 and the water-binding loop of the canonical β-trefoil motif. Although β4’ is missing, a β-strand from elsewhere in the protein packs against β1, effectively acting as surrogate β4’. β2 and β3 are positioned to form a tight hairpin, as in the canonical motif, if not for the presence of an α-helix inserted into the connecting turn.

Although this structure was not identified in our bridging theme analysis, a sequence comparison between the search motif and the βTL motif of IgG-like β-sandwiches reveals a conserved five-residue motif, GQYLA, which located in a structurally equivalent position in both motifs (on β3 and the subsequent loop).

β*-grasp*. Nudix-related hydrolases are a collection of F-groups within the β-grasp evolutionary lineage and comprise H-group 221.4.1. Both a bridging theme and a βTL (**Figure S1**) structural motif **(Figure 3**) were identified in this T-group. Furthermore, although the overall structure of this sequence segment is different in both contexts, β1 of the βTL roughly corresponds to β1 in the canonical β-trefoil motif (**Figure S2**). The phylogenetic distribution of themes in both evolutionary lineages (**Figure S1**) are not consistent with an ancient or persistent evolutionary connection, and instead suggest a more isolated theme sharing event. Nevertheless, this association is interesting because β-trefoils are not associated with catalytic activity, whereas Nudix-related hydrolases, as the name suggests, are enzymes.

Although βTL motifs within the Nudix-related hydrolases are generally characterized by a conserved water molecule between β1 and β2, an exception exists: in NDX2 from *Thermus thermophilus* HB831, the conserved water is excluded by a proline residue, the backbone of which cannot act as a hydrogen bond donor (**Figure S3**). A ‘dry’ β-trefoil motif may therefore be an accessible structural solution, though no such motif could be identified in a β-trefoil. In addition, β-grasp proteins are unique in that the water-binding loop can interact with the conserved water via a serine sidechain, allowing the water molecule to reposition and form four hydrogen bonds (as opposed to the three short hydrogen bonds observed in other protein lineages). Finally, although the core βTL motif present in β-grasp proteins comprises only β1, β2, and the water-binding loop, strands exist on either side of the motif occupying the role of β3 and β4’ of the canonical motif.

## Discussion

### An Updated Protocol Reveals Bridges to an Island Architecture

We, and others^4,7^, have previously analyzed patterns of global sequence fragment sharing across the protein universe, both with^6^ and without^2^ structural constraints. In all cases, connections to the β-trefoil were either marginal or absent. Why, in the present analysis, do we uncover extensive evidence of fragment sharing between β-trefoils and other protein lineages? We attribute the improved sensitivity of our approach to two methodological changes: First, our analysis more directly considers β-trefoil fragments (*i*.*e*., a sliding window of 4 β-strand elements, which is the length of the repetitive motif that forms the β-trefoil architecture; see **Methods** for more details) and not the β-trefoil structure as a whole. When the entire structure – that is, three consecutive β-trefoil motifs – was used, searches were strongly biased in favor of other β-trefoils, thus giving the impression that this evolutionary lineage is an island. This approach was motived both by the accepted models of β-trefoil evolution, in which the β-trefoil emerged from a single oligomerizing β-trefoil motif^9,11^, and previous studies that attempted to uncover associations between β-trefoils and other lineages^12^. Second, we generated HMM profiles from a 70% identity cluster of modern β-trefoils. By using 70% sequence identity clusters to make HMM profiles, we emphasized somewhat more recent evolutionary events than others do (*e*.*g*., Alva and coworkers^7^) and, by virtue of having a finer sampling of sequence space, increase the search space to identify signatures of possible parallel evolution between lineages^8^. This choice is consistent with β-trefoils being a relatively young protein lineage^9,15,17,34^.

For repeat proteins like the β-trefoil, and for protein lineages that lack a highly conserved functional feature, these methodological improvements appear to be crucial to understanding early evolution. We now know that β-trefoils are active participants in sequence fragment sharing across the protein universe, a fundamental shift from the perspective that this fold originated *de novo* and is isolated in sequence space.

### The Origin of β-Trefoils

Experimental fragmentation studies have demonstrated that the β-trefoil architecture can be realized by a trimerizing 32-residue peptide, begging the question: Where did the founding peptide come from in the first place? Is it possible that the β-trefoil is a derived from another protein lineage, perhaps akin to the evolutionary relationship between flavodoxins and TIM barrels^35^? Given that β-trefoils are relatively young, they emerged in a cell already populated by numerous complex proteins, which may be a prerequisite for this evolutionary model. But which evolutionary lineage was most likely to be the parent lineage? Previously studies have suggested potential evolutionary associations with EGF proteins or β-propellers, both of which are observed here. However, on the basis of the analysis presented above, the most probable fragment donor for a founding β-trefoil peptide was the IgG-like β-sandwich, an evolutionary lineage that i) significantly predates the β-trefoil^15–17^ ii) has the strongest signatures of theme sharing with the β-trefoil (**Figure 1**) iii) exhibits theme sharing events with a wide phylogenetic distribution the β-trefoil side and a comparatively narrow phyletic distribution on the IgG-like β-sandwich side (**Figure 2**) and iv) possess a rare example of structural motif convergence within a modern domain (**Figure 3**). Taken together, these observations present the strongest evidence to date for a derived β-trefoil evolutionary model (**Figure 4**), in which (1) the nascent β-trefoil peptide emerged in the context of an IgG-like β-sandwich protein; (2) this peptide ‘budded’ from the parent protein to form a standalone, independently folding trimeric β-trefoil; that was (3) ultimately consolidated onto a single chain by gene duplication and fusion events. Evolution from a fragment, therefore, may be property of a very ancient protein lineage^7^ (*cf*. Rossmanns^8^, P-Loops^36^, and the (HhH)_2_-fold^37^) but may also be a property of a relatively recent protein lineage, like the β-trefoil.

**Figure 4.**
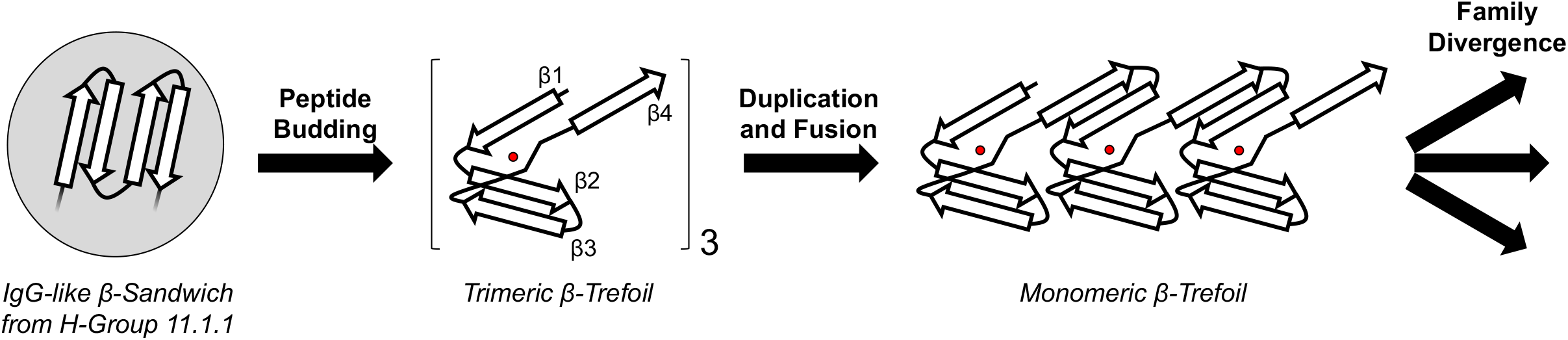
Emergence and evolution of the β-trefoil architecture. The peptide budding model hypothesizes that the first β-trefoil emerged after excision of a segment from an IgG-like β-sandwich (grey shaded circle) to form a trimeric β-trefoil. This model of emergence is suggested by preponderance of bridging themes between IgG-like β-sandwiches and β-trefoils (**Figure 1**), their wide phylogenetic distribution among β-trefoils but narrow distribution within IgG-like β-sandwiches (**Figure 2**), and the observation of a βTL motifs in a modern IgG-like β-sandwich protein (**Figure 3**). The homo-trimeric β-trefoil was then duplicated and fused into a single domain, events predicted by Ponting and Russell^9^ and demonstrated experimentally by Lee and Blaber^11^. The founding monomeric β-trefoil then gave rise to multiple β-trefoil families by both whole domain duplication and *via* a budding-like process of β-trefoil motifs, as described by Broom and coworkers^31^. As drawn, the embedded fragment in the IgG-like β-sandwich does not adopt a β-trefoil motif-like structure and is comprised of four β-strands. However, we do not exclude the possibility that the embedded peptide adopted a β-trefoil-like motif or that it was shorter than a canonical β-trefoil motif and underwent a duplication concurrent with the budding event.

Finally, we note that the budding of a peptide from a folded domain to found a new protein family has been observed before: Broom and coworkers^31^ argued that a single β-trefoil motif from a monomeric β-trefoil protein can be duplicated and fused to yield a new family of β-trefoils, a process of modular evolution. Likewise, the helix-hairpin-helix (HhH) motif is known to exist as both an embedded functional element and as a standalone domain with a two-fold axis of rotational symmetry, demonstrating that transitions between embedded structural elements and independently folding symmetric domains occurs elsewhere in the protein universe^7,38^.

## Conclusions

Modern proteins are a web of evolutionary associations that manifest across various levels of organization, from genes to domains to sequence segments. Advancements in search algorithms and analysis protocols, such as the bridging themes presented here, will undoubtably uncover even more associations between globally unrelated evolutionary lineages – revealing that seemingly isolated folds, like the β-trefoil, are in fact active participants in fragment sharing. Characterizing the presence and extent of bridging themes between evolutionary lineages can provide valuable insight into protein domain emergence^7,8^; in the present case, by suggesting that the β-trefoil is a derived fold that arose from an IgG-like β-sandwich.

## Data Availability Statement

All data used in this manuscript is available either as a supplemental file or in a publicly accessible database.

## Acknowledgements

This work was supported by NSF award #1724300 (S.E.M.). We gratefully acknowledge Dr. Michael Blaber and Dr. Dragana Despotović for thoughtful discussions during the preparation of this manuscript.

## Notes

### Competing Interest Statement

The authors have declared no competing interest.

