## Supplemental Information for "Evidence for the Emergence of β-Trefoils by ‘Peptide Budding’ from an IgG-like β-Sandwich"

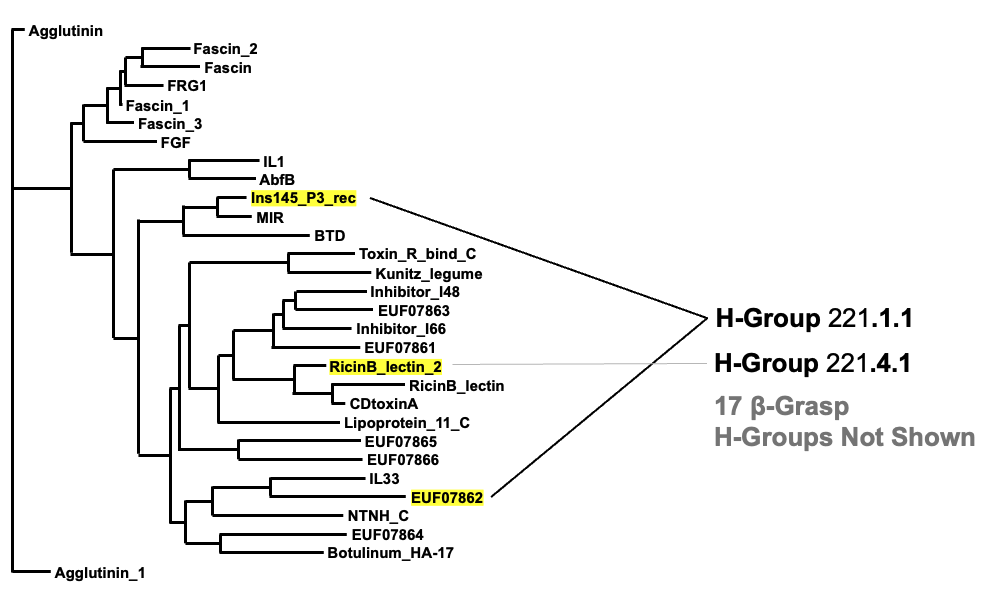

Figure S1. The distribution of β-grasp bridging themes across the β-trefoil phylogenetic tree.

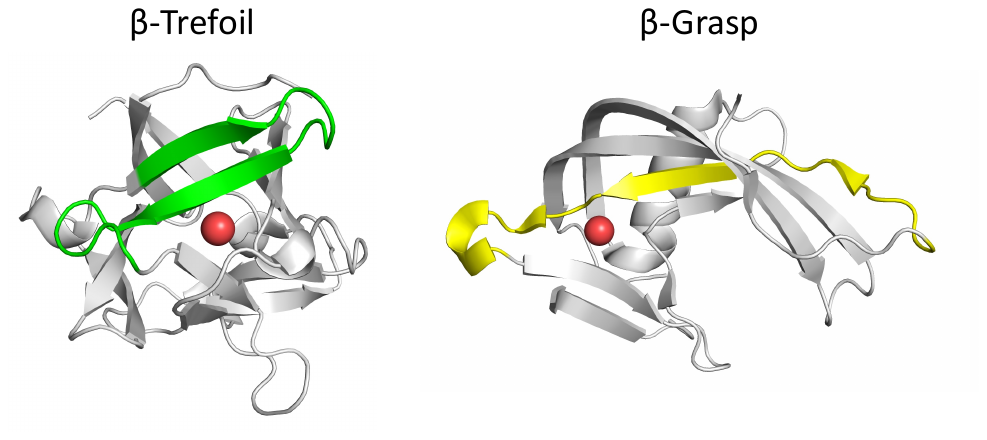

Figure S2. An overlapping βTM and bridging theme. Regions of the domains associated with the bridging themes are colored green or yellow. The conserved water molecule of the βTL motif I shown as a red sphere. ECOD domains are e2vseA2 (β-trefoil F-group RicinB_lectin_2) and e2azwA1 (β-grasp H-group 221.4.1).

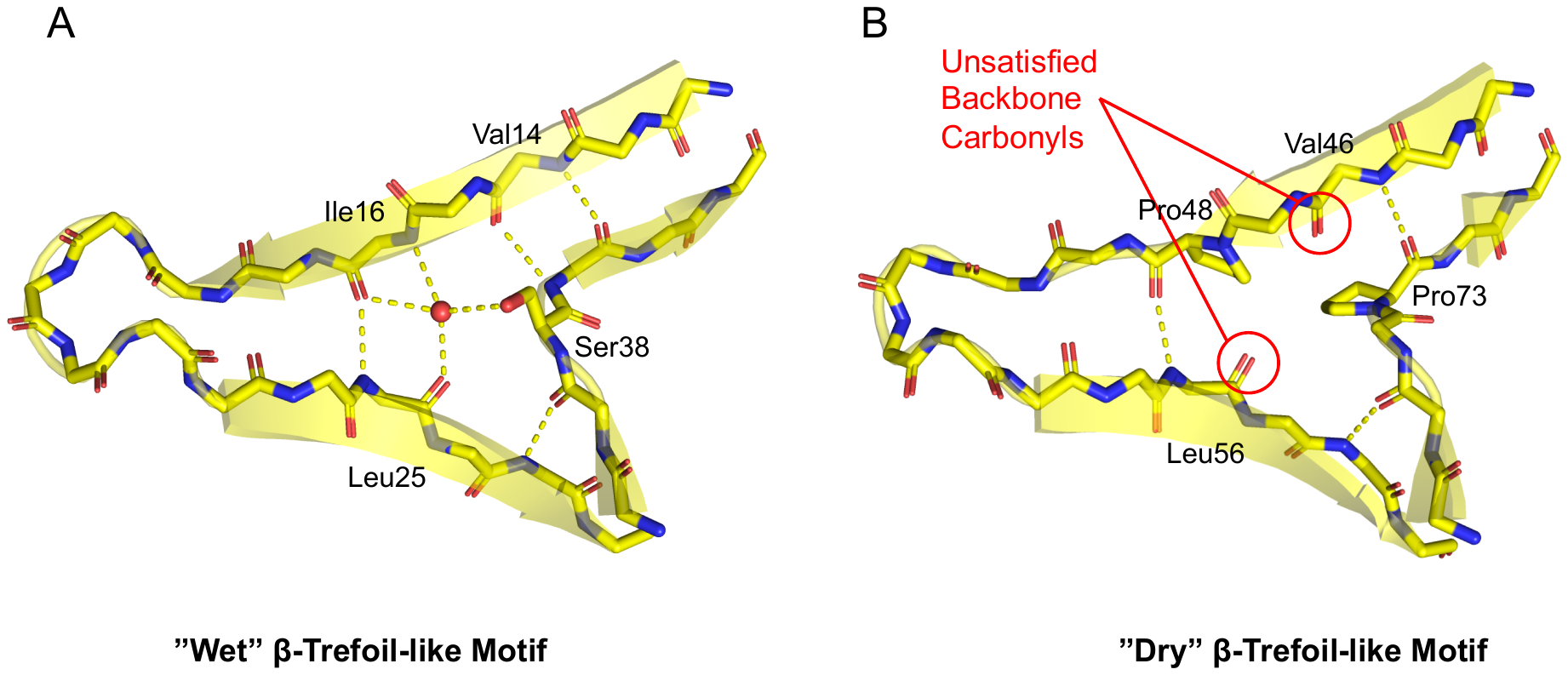

**Figure S3**. β-trefoil-like motif without a conserved water in the β-grasp evolutionary lineage. **A**. In the canonical β-trefoil motif, a conserved water bridges β1 and β2 (red sphere), a feature that is retained in many Nudix hydrolases. Shown here is ECOD domain e2o1cB1 (F-group 221.4.1.3). **B**. In some Nudix hydrolases, however, a proline residue on β2 of the βTL motif precludes water binding. In ECOD domain e2yvpA2 (also F-group 221.4.1.3), formation of a ‘dry’ β-trefoil motif, the result of two proline residues – one on β1 and one on the loop just after β2 – results in two unsatisfied backbone carbonyls (annotated with red circles).
